# Accelerating Maximal-Exact-Match Seeding with Enumerated Radix Trees

**DOI:** 10.1101/2020.03.23.003897

**Authors:** Arun Subramaniyan, Jack Wadden, Kush Goliya, Nathan Ozog, Xiao Wu, Satish Narayanasamy, David Blaauw, Reetuparna Das

## Abstract

**Motivation:** Read alignment is a time-consuming step in genome sequence analysis. In the read alignment software BWA-MEM and the recently published faster version BWA-MEM2, the seeding step is a major bottleneck, for instance, contributing 38% to the overall execution time in BWA-MEM2 when aligning single-end whole human genome reads from the Platinum Genomes dataset. This is because both BWA-MEM and BWA-MEM2 use a compressed index structure called the FMD-Index, which results in high memory bandwidth requirements for seeding, primarily due to its character-by-character processing of reads.

**Results:** We propose a memory bandwidth-aware data structure for maximal-exact-match seeding called Enumerated Radix Tree (ERT). ERT trades off memory capacity to improve seeding performance (∼60 GB index for human genome). Together with optimizations to the seeding algorithm and mate-rescue step, ERT when integrated into BWA-MEM2 speeds up overall read alignment by 1.28× and provides up to 2.1× higher seeding performance while guaranteeing identical output to the original software. Furthermore, we prototype an FPGA implementation of ERT on Amazon EC2 F1 cloud and observe 1.6× higher seeding throughput over a 48-thread optimized CPU-ERT implementation.

**Availability and implementation:** https://github.com/arun-sub/bwa-mem2

## 1 INTRODUCTION

Read alignment is one of the major compute bottlenecks in secondary analysis. Every read needs to be *aligned* to a position in the reference genome. The seed-and-extend heuristic (Baeza-Yates and Perleberg, 1996) is commonly used for read alignment. Seeding finds a set of candidate locations (*hits*) in the reference genome where a read can potentially align. Hits for a read are determined by finding exact matches for its substrings (*seeds*) in the reference. The seed extension phase then uses approximate string matching to select the hit with the best score as the read’s alignment position.

The seeding step is a major bottleneck in read alignment software such as BWA-MEM. For instance, seeding contributes ∼38% to the overall run time of BWA-MEM2 (Vasimuddin *et al.*, 2019). In addition to read alignment, seeding is also an important kernel in several other applications: metagenomics classification (e.g., Centrifuge (Kim *et al.*, 2016)), de-novo assembly (Simpson and Durbin, 2012)) and read error correction (Greenstein *et al.*, 2015). Therefore, there is a need for fast and efficient solutions for seeding.

The primary performance bottleneck in seeding is memory bandwidth resulting from irregular memory accesses to the FMD-index. The highly compressed FMD-index in BWA-MEM (4.3 GB for human genome) trades off high memory bandwidth for small memory space. However, memory bandwidth is a precious resource in modern processors that has not scaled proportionately with core counts (Wulf and McKee, 1995). We find that on real whole human genome reads, ∼40% CPU cycles are spent waiting for data from memory elements such as on-chip caches and DRAM. Furthermore, irregular memory accesses in seeding cannot take full advantage of the memory system which has been optimized for regular accesses with small stride (i.e., accesses with high spatial locality). As a result, seeding only utilizes 11.7% of peak DRAM bandwidth in a 48-thread multicore system.

In contrast, we propose a data structure for seeding that makes the opposite trade-off: it trades off increased memory space for reducing bandwidth required, while still fitting within a modern server’s main memory (64 GB) as shown in Figure 1. This design tradeoff is similar in spirit to that made in BWA-MEM2 (which uses a lower compression factor for the FMD-index, resulting in a 42 GB index for the human genome), but our solution further improves bandwidth efficiency by virtue of supporting multi-character lookup and exploiting reuse opportunities by redesigning the seeding algorithm.

**Fig. 1.**
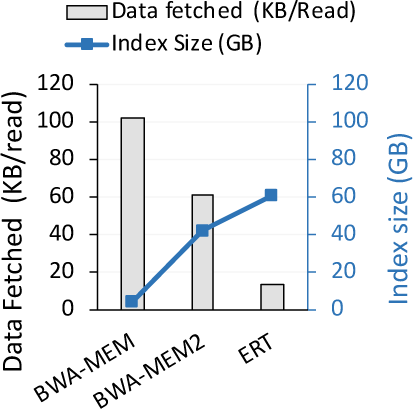
Trade-off between index size and data required for seeding.

We refer to our bandwidth-efficient data structure as Enumerated Radix Trees (ERT). Like FMD-index, ERT enables variable length exact match search functionality. But, unlike FMD-index, it avoids iterative lookup for every base-pair on a large structure which can result in poor performance. It achieves this by coalescing all substrings in a reference genome that start with the same k-mer together (where k is less than the minimum length for a seed), and representing them using a variant of a radix tree. As we discuss later, ERT allows *multiple consecutive base-pairs to be matched with one lookup*, and exhibits better spatial locality than the FMD-index. ERT also helps *reduce computation when substrings* within a read that need to be matched with the reference *overlap* using a prefix-encoded radix tree. ERT increases data efficiency of seeding (data fetched per read) by 4.3× as shown in Figure 1 and improves overall seeding performance of BWA-MEM2 by 1.6–2.1×.

## 2 BACKGROUND AND MOTIVATION

### 2.0.1 Seeding Algorithm in BWA-MEM

The seeding algorithm in BWA-MEM is based on identifying substrings that have super-maximal exact matches (SMEMs) with the reference genome (Li, 2013) as shown in (Figure 2(a)). A maximal exact match (MEM) is an exact match that cannot be extended in either direction in the read. An SMEM is a maximal length match (MEM) that is not *fully contained* in any other MEM. Figure 2 (b) shows the steps involved in determining SMEMs for a sample read and reference pair.

**Fig. 2.**
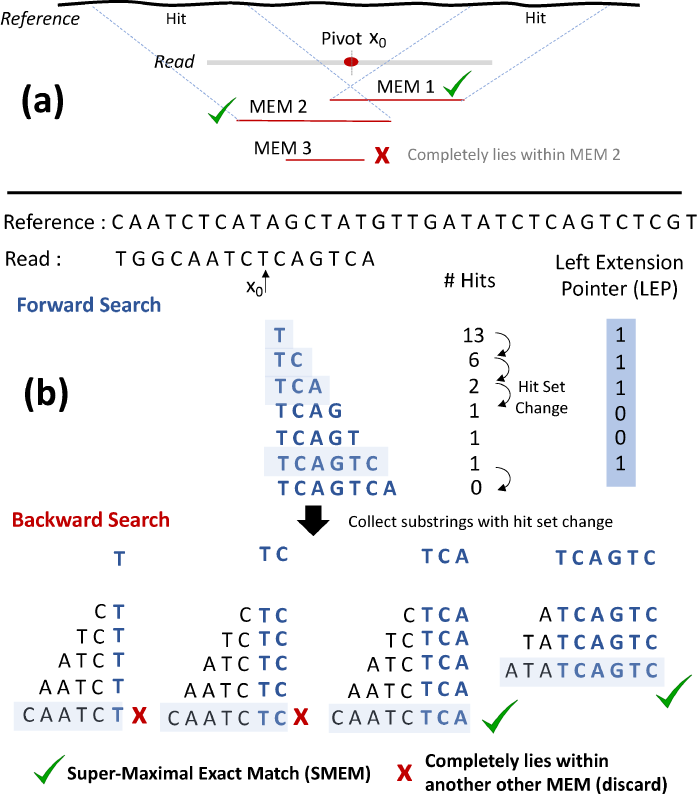
(a) Super-Maximal Exact Matches example. (b) Forward and backward search to identify super-maximal exact matches (SMEMs).

To identify SMEMs and their locations in the reference genome, both BWA-MEM and BWA-MEM2 use a compressed data structure called the FMD-index (Ferragina and Manzini, 2000; Li, 2012) which is built using both strands of DNA (∼6 billion characters of the human genome). The FMD-index consists of: (1) the *suffix array (SA)*, which contains the locations of lexicographically sorted suffixes of the reference genome `R`, (2) the *Burrows Wheeler Transform (BWT)*, computed as the last column of the cyclically sorted suffixes of the reference, (3) the *count table (C)* which stores the number of characters in `R` lexicographically smaller than a given character `c` and (4) the *occurrence table (Occ)* which stores the number of occurrences of a character up to a certain index in the suffix array. Using the FMD-index, SMEMs are identified in two steps:

#### (1) Forward search

For a given query position in the read (e.g., pivot *x*_0_ in Figure 2), subsequent base pairs to its right are looked up one at a time in the FMD-index to find the longest exact match in the forward direction. The position of mismatch becomes the start of the next pivot and the process repeats. During forward search from a pivot, all the positions in the read where there is a *change* in the set of candidate reference locations (hits) are marked (left extension points (LEP) in Figure 2). Only these positions are used as the starting query positions to identify potentially longer exact-matches extending in the backward direction. Other positions are guaranteed to produce MEMs that are contained within those identified from LEP.

#### (2) Backward search

For each query position identified in the previous step (LEP), subsequent bases to the left of the pivot are looked up one at a time to find the longest exact match in the backward direction. After this process, SMEMs are identified by discarding MEMs fully contained in other longer matches. The locations of these SMEMs in the reference genome (hits) are then determined by a suffix array lookup (can take multiple occurrence table lookups in BWA-MEM, since the suffix array is sampled) and passed on to the seed-extension stage. SMEMs obtained during seeding are assumed to be part of the final alignment.

### 2.0.2 FMD-Index Seeding Bottlenecks

The FMD-index allows the lookup of query `Q` of length `N` in reference `R` using approximately 𝒪(*N*) memory operations. Detailed descriptions of the FMD-index can be found in (Ferragina and Manzini, 2000; Li, 2012). Like BWA-MEM, BWA-MEM2 also uses the FMD-index for seeding, but uses a lower compression factor in its implementation to reduce memory bandwidth requirements. In particular, the occurrence table used for performing range queries on the FMD-index is decompressed by 4× and the suffix array to identify locations of substrings in the reference genome is fully decompressed. These changes increase the FMD-index size to 42 GB (12 GB occurrence table + 30 GB suffix array) (Vasimuddin *et al.*, 2019) compared to 4.3 GB in BWA-MEM. Starting from a single character in the read, the FMD-index enables forward and backward MEM searches to determine the number of hits of progressively longer substrings using at most 2 extra memory lookups per character.

Software implementations of the FMD-index (e.g., BWA-MEM) have attempted to improve the spatial locality of FMD-index lookups in two ways: *First*, occurrence table entries are typically co-located with portions of the BWT in tightly packed cache-line aligned (i.e., 64 B) data structures to reduce accesses to main memory (can take 2–50× longer than a cache access). *Second*, backward search passes for substrings sharing the same prefix (e.g., `TCAGTC` and `TCA` in Figure 2) are performed in lock-step leading to access of FMD-index data belonging to the same or nearby cache lines (Li, 2013; Xin *et al.*, 2015). However, the initial few memory lookups touch different parts of a ∼42 GB data structure and rarely exhibit a regular, predictable memory access pattern. This reduces the effectiveness of multi-level caching in modern processors and imposes high memory bandwidth requirement.

## 3 METHODS

### 3.1 Enumerated Radix Tree Index Design

In this section, we present *Enumerated Radix Trees (ERT)*, which is built from the ground up to overcome the low data efficiency of FMD-index based seeding. Like the FMD-index, to support both forward and backward searches, ERT is built for the concatenated reference: consisting of both the forward and reverse complemented strands of DNA.

#### 3.1.1 K-mer Index

FMD-index stores a compressed representation of the set of *all suffixes that exist in the reference genome* in lexicographical order. We now consider a substring of length k in the read (referred to as a *k-mer*). Due to natural genome variation and machine read error, not all k-mers will exist in the reference and, hence, in the FMD-index. Therefore, when looking up a k-mer in the FMD-index, we must start with a 1-mer and grow the string, character by character, for as long as it exists in the FMD-index, or till we reach the desired k-mer length. This iterative, character-by-character access to the FMD-index substantially increases the required number of DRAM accesses, creating a memory bottleneck. This is further aggravated by the fact that accesses to the index rarely follow lexicographic order, making it difficult to exploit locality over such a large window (i.e., set of all suffixes of the k-mer).

To overcome these two limitations, we instead enumerate *all possible k-mers* (whether they exist in the reference or not) and store them in an index table. Prior work (Xin *et al.*, 2015) also made a similar observation to increase locality in the first few steps of FM-index search. For each k-mer (an index entry), we then store all its suffixes in the reference. Since all possible k-mers are represented in the index, *k characters* from the read can be looked up in a single memory access, significantly reducing the number of DRAM accesses. Furthermore, subsequent accesses to the suffixes of the k-mer have much improved spatial locality, since they are co-located together. To capture intermediate results from each of the k single character lookups, we leverage memoization, wherein we precompute and store the results of looking up every possible k-character permutation in a k-mer index table (LEP in Figure 2 b.). Figure 3 shows an example index table enumerating all 6-character substrings.

**Fig. 3.**
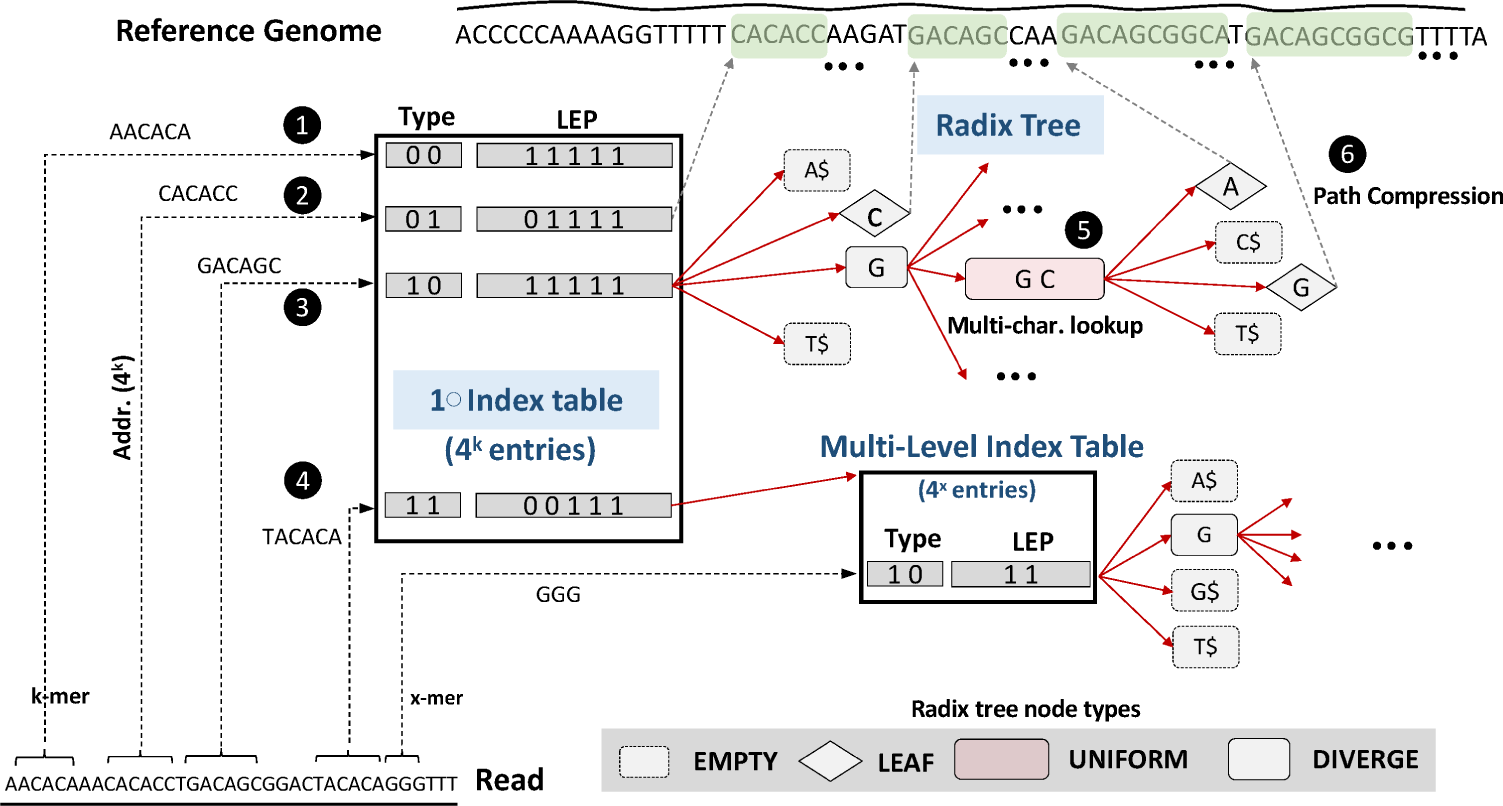
Enumerated radix tree (ERT) example. ERT supports multi-character lookup using a multi-level index table and radix tree. Each index table entry contains a `Type` field, k-1 bit `LEP` field (Section 2) and a `pointer` field that indicates the address of the root node of a radix tree (not shown). `Type` can be any of the following: (1) `EMPTY`: k-mer is absent in the reference. (2) `LEAF`: k-mer is unique (i.e., has the same suffix at all occurrences in the reference). (3) k-mer exists in the reference and has a pointer to the root of the radix tree. (4) k-mer has very high number of unique suffix strings (T > 256) in the reference. The index table entry for this k-mer points to a 2nd-level index table to succinctly represent these suffix strings. ERT represents a singleton path using a variable-size internal node (`UNIFORM`) supporting multi-character lookup (5). If a singleton path ends in a leaf, it is truncated at its start with a `LEAF` node that points to the reference genome (path compression, 6). ERT also includes `EMPTY` nodes (ending with $) to indicate absence of prefixes in the reference and `DIVERGE` nodes which have more than one valid child branch paths for the prefix.

To **choose k**, we observe that BWA-MEM only reports SMEMs greater than a certain minimum length (e.g., 19). We choose k=15 to keep the size of index table tractable (*O*(4^*k*^)), i.e, 1 G entries when *k* = 15. Later, in section 3.5.1 we discuss a solution to effectively increase *k* by selectively using a multi-level index.

#### 3.1.2 Customized Radix Tree

The next question is how to store the suffixes of a k-mer in an index entry, so that we can support MEM searches for strings longer than `k`. One option is to augment the index table with an FMD-index, and iteratively grow the k-mer prefix. However, even within the subset of all suffixes sharing the same k-mer prefix, FMD-index lookups have poor locality. Also, they still operate with a single character at a time.

To overcome this problem, we observe that a radix tree can naturally support *multi-character lookups*. This is because in a radix tree, we can merge all singleton paths into a single node, thereby addressing a multiple character lookup with a single memory access. Figure 3 shows a radix tree for one k-mer in the index table (note radix is 4 for the genome alphabet). The proposed ERT merges singleton paths (*GC* in Figure 3) using variable-size internal nodes that store the full singleton path string (designated as UNIFORM). A singleton path is encountered when all paths in the tree from a certain node onward share a common string.

##### Early Path Compression

To further improve the space-efficiency of the ERT, we observe that a k-mer frequently becomes *unique* in the reference genome as it increases in length. This means that, past a certain length, a prefix is followed by a single, unique suffix string in the reference genome. This would introduce a UNIFORM node in the ERT with a singleton string of characters (up to the length of the read). To avoid storing this long string, we instead replace it with a pointer to the occurrence of this string in the reference genome. In Figure 3 we show how in the ERT, these nodes are marked as *leaf nodes*, containing a single pointer. Leaf nodes encountered during a MEM search are decompressed, by fetching the full reference string corresponding to the reference pointer stored at the leaf node. Note that the pointer in the leaf node is required regardless of this compression technique since it is necessary to indicate the location of the traversed k-mer in the reference genome. Hence, it does not present any storage overhead. Instead, this optimization results in ∼2× space savings and was critical for being able to store the full human genome in under 64GB of storage, which is a common configuration for servers.

### 3.2 Index Construction and Operation

The ERT `k-mer` index table and corresponding radix trees are built by first enumerating all possible k-mers and then querying the FMD-index to grow the trees according to all existing sequences in the reference. Each k-mer and ERT path corresponds to a unique sequence in the reference. The locations of these sequences are stored as pointers at the leaves of the tree, as noted above. Note that if a particular k-mer does not exist (referred to as *EMPTY* in Figure 3), we do not store a pointer to an ERT tree. In our implementation, for the human genome, 38.8% of the index entries are empty when *k* = 15. For an empty entry, we still compute its LEPs and store it in the index table to indicate at which positions along this k-mer a backward traversal must be initiated. ERT construction is multi-threaded and takes < 1 hr wall-clock time, ∼32 GB peak memory for the human genome on a 48-thread CPU (m5d.12xlarge).

Once constructed, we can use the ERT to search for MEMs according to the SMEM seeding algorithm (Section 2). For a given `k-mer` scanned from a read, based on the starting position of the MEM, we do the following: **(1)** The index table is looked up using the `k-mer` with a single DRAM access. If an entry in the index table exists, the root of the `k-mer` tree is also fetched with a second memory access. **(2)** The branches in the `k-mer` tree are then traversed according to the remaining base pairs in the read until a leaf node is encountered or a dead end is reached (i.e. no further characters in the read match with strings existing in the reference). **(3)** If a leaf node is reached, the reference sequence corresponding to that leaf is fetched with a DRAM access to determine the final characters matching with the read. **(4)** If we reach a dead end in the tree, we have found the end of the MEM. At this point all locations where this MEM exists in the reference (i.e., all leaf nodes in the downstream sub-tree) are gathered using a depth-first traversal, referred to as *leaf gathering*. **(5)** Each time the path in the ERT traverses a node with divergence, an LEP is marked since the divergence indicates that the number of hits is divided across the divergent paths from that node and is decreasing. **(6)** If the forward search reaches a dead end (or the end of the read), a backward traversal is instigated for each LEP position identified. The backward traversal operates in the same way as the forward traversal and uses the same ERT data structure but instead operates on the reverse complemented read. Note that base-pairs A and T and base-pairs C and G are complements of each other.

### 3.3 Data Structure Optimization: Prefix-Merged Radix Tree

The goal of prefix-merged radix trees is to re-use work across MEM searches from consecutive positions in the read.

#### Opportunity

In the seeding computation, the time spent doing backward MEM searches is ∼2× that of forward search, making it important to optimize this step. On average, we find that there are ∼10 backward searches for each forward search from a pivot. Also, it is common to observe backward searches from adjacent query positions in the read (consecutive bits of LEP are ‘1’). Normally, these lead to multiple independent index table lookups and tree traversals as shown in Figure 4.

**Fig. 4.**
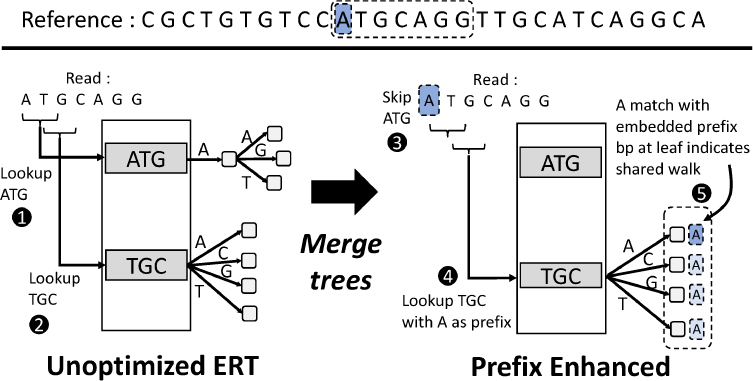
Merging radix-trees by adding prefix data at the leaf nodes allows ERT to leverage prefix information to perform multiple MEM searches in a single tree traversal.

#### Insight

In the unoptimized ERT, there exists a radix tree for each k-mer that occurs in the reference, including adjacent, sliding window k-mers (e.g., `ATG` and `TGC`). We recognize that radix trees for adjacent k-mers contain redundant information and that the information contained in one of the trees can be reconstructed from the adjacent k-mer’s tree by storing prefix information at each of its nodes. In the example shown in Figure 4, a string `ATGC`, which is normally found by accessing the `ATG` tree can be instead reconstructed from the `TGC` tree by indicating the presence or absence of prefix character `A` in each of the nodes of `TGC`’s tree.

The key observation is that with such a prefix-merged radix tree, multiple backward searches (`TGCxyz` and `ATGCxyz`) can be performed in a *single index table lookup and tree traversal* by checking for prefix character matches at each visited node. In Figure 4, when we reach the leaf node represented by string `TGCA`, we can also match character `A` from the read as prefix, resulting in the MEM represented as `ATGCA`. This reduces two backward extensions into one.

#### Design

Augmenting each of the nodes with prefix information in order to merge k-mer trees takes up significant space and offsets the benefit from merging trees. Therefore, in our prefix optimized ERT, only leaf nodes are augmented with prefix characters (2 bits per prefix character) found at the corresponding reference positions (Figure 4). Storing prefix information at the leaf nodes is sufficient as prefix information at each of the internal nodes can be reconstructed by visiting all of the leaf nodes in its corresponding sub-tree. If any of the leaf nodes of an internal node’s sub-tree contains the desired prefix character, then the internal node also contains the prefix character. While storing prefix information at internal nodes does have the benefit of terminating some backward searches early in case of prefix mismatch, we found that the space overhead outweighed the performance benefits.

Another design choice to be made for prefix-merged ERT is the choice of prefix length. We observe that each backward search on average matches ∼1 prefix character at the leaf nodes, resulting in 50% fewer backward searches. As a result ERT supports 1-character prefix at leaf nodes. Although the above discusses the benefits of prefix-merged radix trees in the context of backward searches, it must be noted that forward MEM searches can also benefit from this optimization when initiated from adjacent positions in the read.1

### 3.4 Algorithm Optimization: Pruning Wasteful Backward Searches with Zigzag Seeding

After performing forward search from the pivot (Figure 5 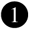), we obtain the LEP vector 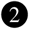 to guide backward search 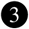. Typically backward search is performed starting from each query position in the read where the set of candidate hits changes (as given by set bits in the LEP vector). In the original seeding algorithm, backward search proceeds in the right-to-left order for each bit set in the LEP vector, starting from the longest match (rightmost ‘1’ bit in LEP) and ending at the shortest match (leftmost ‘1’ bit in LEP). However, as can be seen in Figure 5 (a), many of these backward extensions end in MEMs that are fully contained in previously identified SMEMs. These MEMs are discarded by the seeding algorithm.

**Fig. 5.**
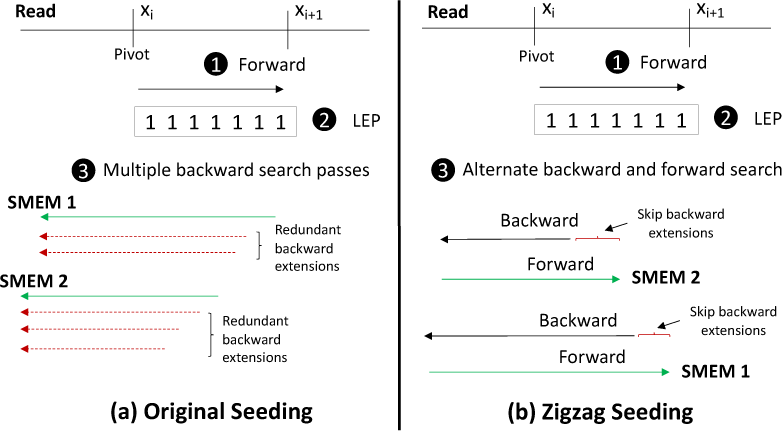
(a) Original seeding algorithm that performs multiple backward search passes sequentially leading to redundant backward extensions. (b) Redesigned seeding algorithm that interleaves backward and forward searches to skip redundant backward extensions.

Performing backward extensions for LEP positions that do not lead to SMEMs is wasteful. Ideally, we would like to perform only those backward extensions that lead to non-overlapping MEMs. To achieve this, we redesign the seeding algorithm to alternate between forward and backward search in a *zigzag* fashion as shown in Figure 5 (b). Instead of starting backward search at the rightmost set LEP position, we start backward search at the pivot and extend leftward until no longer match can be found. We later extend the same match beyond the pivot in the forward direction until no longer match can be found. Backward searches from LEP positions in the read that lie within the forward match can safely be skipped since they are guaranteed to produce shorter fully-overlapping matches. The interleaved backward-forward search is repeated from the next set LEP position beyond the forward match as shown in Figure 5 (b).

### 3.5 Space Reduction

#### 3.5.1 Multi-Level Index

##### Opportunity

Enumerating all k-character prefixes in the index table can have prohibitive space overheads for large k. For example, 19-mer table has 4^19^ entries, resulting in 2 TB of space, assuming 8 bytes per entry. However, the human genome is not a random string of characters from the genome alphabet. The repetitive nature of the human genome makes the distribution of hits (or leaf nodes in the radix tree) for different k-mers heavily skewed.

##### Insight

We leverage the skewed distribution of k-mers in the human genome to design a multi-level index table. For a given number of hits `X`, Figure 6 shows the number of k-mers in the human genome that have hits > `X`. It can be seen that very few k-mers (∼ 0.01%) have greater than 1000 hits. However, these k-mers have dense radix trees, which can be compactly represented using an index table as shown in Figure 6.

**Fig. 6.**
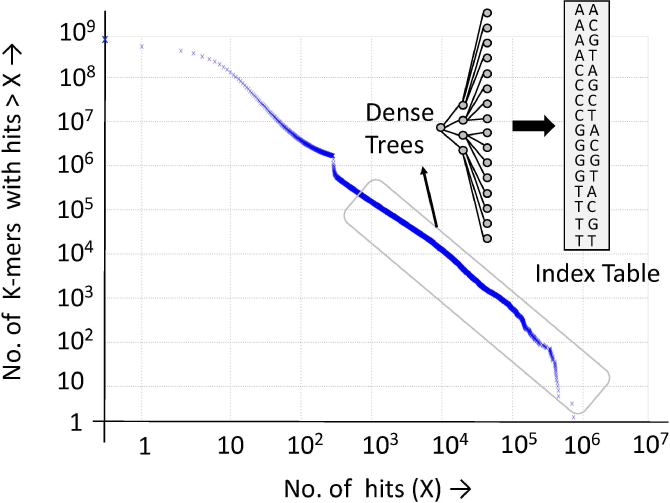
Figure showing skewed hit distribution for k-mers. The few k-mers with a large number of hits (dense trees) are represented with an index table.

##### Design

Instead of enumerating all k-character prefixes for large k, we decompose the index table into two levels (Figure 6), wherein the first level enumerates all k-mers and the subsequent level enumerates all *x*-character suffixes for a subset of k-mers (such that *k* + *x* = min. SMEM length). The multi-level index table further extends the benefit of multi-character lookup. Another way to visualize the multi-level index is as a high fan-out tree, with the root being the k-mer and the children being all *x*-character suffixes for the k-mer. While choosing a larger *x* helps reduce tree traversal time, for the human genome we were able to accommodate up to *x* = 4 (fan-out = 256) for a subset of 15-mer dense trees without increasing space overheads (only 0.35 % of all 15-mers > 100 leaf nodes). Compared to *x* = 1, *x* = 4 improves CPU performance by 10%. Since most trees are shallow (83% of leaf nodes have depths <= 8), we did not explore more than two-levels or high fan-out for internal nodes of ERT.

#### 3.5.2 Variable-Size Radix-Tree Nodes

Radix-tree nodes also contribute significantly to the size of the ERT index. Each radix tree node in ERT is variable-size consisting of the following: (1) 2-bit type (i.e., EMPTY, LEAF, UNIFORM and DIVERGE) for each of its children nodes, (2) *n*-bit pointers to children nodes, (3) *m*-character string to represent singleton paths and (4) *log*_2_|2*R*|-bit pointers to the reference genome (length = |*R*|) in case of LEAF nodes. Since few k-mers have large trees, there exists opportunity to reduce the bit-width of pointers to children nodes for many k-mers. We choose between 16, 24 or 32-bit pointers for each k-mer during index construction based on the size of its tree. For the human genome, >99% k-mers use 16-bit pointers.

### 3.6 Improving Seeding Locality with K-mer Reuse

Given the highly repetitive nature of the human genome and high coverage of sequenced reads needed to overcome sequencing errors (each position in the reference genome can be covered by 30-50 reads on average), few unique k-mers tend to appear frequently in a batch of reads. As a result, there exists opportunity to save memory bandwidth by preventing multiple radix tree fetches for the same k-mer. Unfortunately several radix trees need to be accessed to find seeds for a read, and their aggregate size exceeds that of on-chip caches. As a result, in the baseline, a radix tree usually gets evicted before it can be reused by the same k-mer appearing in a different read. To mitigate this problem, we first perform forward search for a batch of reads, identify all the unique k-mers that are to be used in backward search, fetch each radix tree once for each unique k-mer and perform all backward searches involving that k-mer’s tree before moving to the next k-mer. We refer to this technique as *k-mer reuse*. Supplementary Material includes more details on *k-mer reuse* hardware implementation and methods to further improve spatial locality of seeding.

### 3.7 FPGA Prototype

While seeding is inherently a memory-bound algorithm, CPU implementations can only issue limited number of parallel memory requests and hence cannot saturate memory bandwidth (only uses 11.5% of peak bandwidth). Current GPUs are not well-suited because of significant memory divergence during tree traversal (Wang *et al.*, 2019). To make better use of available memory bandwidth, we design a custom seeding accelerator and prototype it on an FPGA.

The seeding accelerator architecture is shown in Figure 7. It is composed of multiple parallel seeding machines connected to the available DRAM channels using a crossbar network. Each seeding machine is composed of a control processor that issues commands to three types of processing elements. Each processing element performs a sub-task associated with SMEM identification (i.e. index table lookups, walking ERTs, and DFS leaf gathering). When a processing element issues a memory request to the Data Fetcher–a rudimentary address generation unit and memory controller– and a memory stall occurs, the processing element immediately switches to a new context. This context switching greatly increases compute density of each seeding machine and is essential to an FPGA implementation with limited logic and routing resources.

**Fig. 7.**
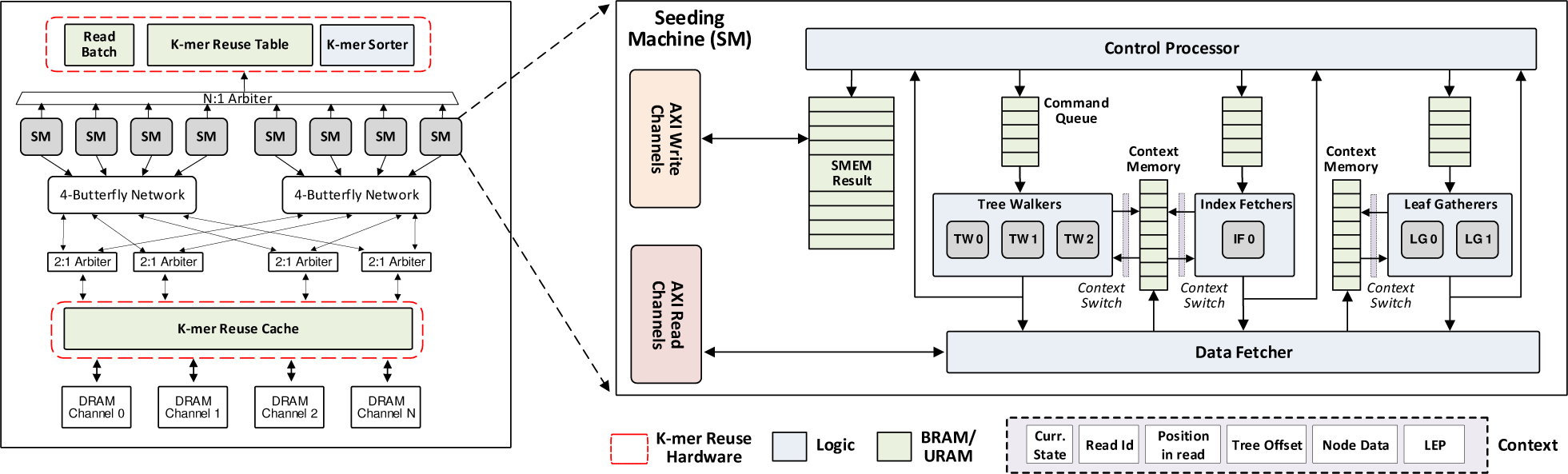
Accelerator architecture. Each Tree Walker (TW) is responsible for scanning a read, walking ERT Trees, and computing candidate SMEMs. Each Tree Walker can switch between multiple contexts to help hide memory latency. The Data Fetcher (DF) is responsible for serving ERT and reference fetch requests to DRAM. The Control Processor (CP) coordinates read fetch, and k-mer reuse phases. The K-mer reuse table stores the results of the K-mer sorter such that after forward extension, identical k-mers used in backward extension are stored together.

We prototyped and verified our seeding accelerator on Amazon’s EC2 F1 FPGA cloud environment. We chose the `f1.4xlarge` instance with 2 FPGAs and equivalent bandwidth as the CPU configuration (68 GB/s peak bandwidth per FPGA). Each FPGA in the F1 instance is a Xilinx XCVU9P with 2,586K logic cells, 36.1 Mbits of Block RAM and 270 Mbits of UltraRAM. System configuration and synthesis results are shown in Table 1.

**Table 1.** FPGA Configuration and Synthesis Results per Xilinx Virtex Ultrascale+ (VU9P) FPGA.

| Component | Configuration | LUT (%) | BRAM (%) | URAM (%) |
| --- | --- | --- | --- | --- |
| Index FU | 1 x 8 | 0.32 | 0 | 0 |
| Walker FU | 3 x 8 | 13.76 | 0 | 0 |
| Leaf Gathering FU | 2 x 8 | 3.36 | 0 | 0 |
| Command Queues | 0.72 KB x 8 | 1.92 | 6.08 | 0 |
| Context Memories | 17.6 KB x 8 | 0 | 15.04 | 3.28 |
| Control Processors | 1 x 8 | 0.56 | 0 | 0 |
| Data Fetcher | 1 x 8 | 3.68 | 0 | 0 |
| SMEM Result Buffer | 2.3 KB x 8 | 0 | 0 | 13.28 |
| MISC. |  | 1.12 | 0 | 0 |
| **Seeding Machines Total** | **1 x 8** | **24.72** | **21.12** | **16.64** |
| **K-mer Sorter** | — | 1.95 | 0.3 | 26.77 |
| **K-mer Reuse Cache** | 4.01 MB | 10.04 | 5 | 18.33 |
| **Seeding Accelerator Total** | **1** | **37.22** | **26.41** | **61.77** |

## 4 RESULTS

We used BWA-MEM v0.7.17 (Li, 2013) and BWA-MEM2 (Vasimuddin *et al.*, 2019) v2.0.pre2 as baselines for comparison. We replace FMD-index seeding in BWA-MEM2 with ERT-based seeding. We also redesign the mate-rescue step in BWA-MEM2 to reduce time spent in sorting. Human whole-genome paired-end short reads (read-lengths 100–151 bp) from Illumina Platinum Genomes (Eberle *et al.*, 2017), Illumina Public Data downloaded from BaseSpaceHub (Project HiSeq 2000 Tumor Normal WGS – `HCC1187BL, HCC2218BL`. Project Polaris 1 Diversity Cohort – `HG03521, HG01624, HG00613`) and 1000 Genomes Project Phase 3 (`ERR3239`) are aligned against the Homo sapiens assembly38 reference. We used `m5d.12xlarge` (48-thread, 192 GB RAM) instance for Illumina Platinum Genomes as some of those (e.g., `ERR194146`) required ∼135 GB peak memory with ERT. For the other datasets, we use the `m5d.8xlarge` (32-thread, 128 GB RAM) instance. Our FPGA prototype implementation uses the `f1.4xlarge` instance as described in Section 3.7. For the software version of ERT, we omitted the k-mer reuse optimization because it requires a software managed cache which slows down the CPU implementation. We verified that ERT produces identical output as BWA-MEM for all the datasets evaluated.

### 4.1 Overall Read Alignment Performance

Table 2 shows the overall wall-clock time for the different datasets. It includes both index loading and read alignment time. On average, ERT improves read alignment performance of BWA-MEM by 2.59× and BWA-MEM2 by 1.28×. ERT provides greater benefits when seeding time is dominant as can be seen for the `HCC1187BL,` `4.2 HCC2218BL` datasets.

**Table 2.**
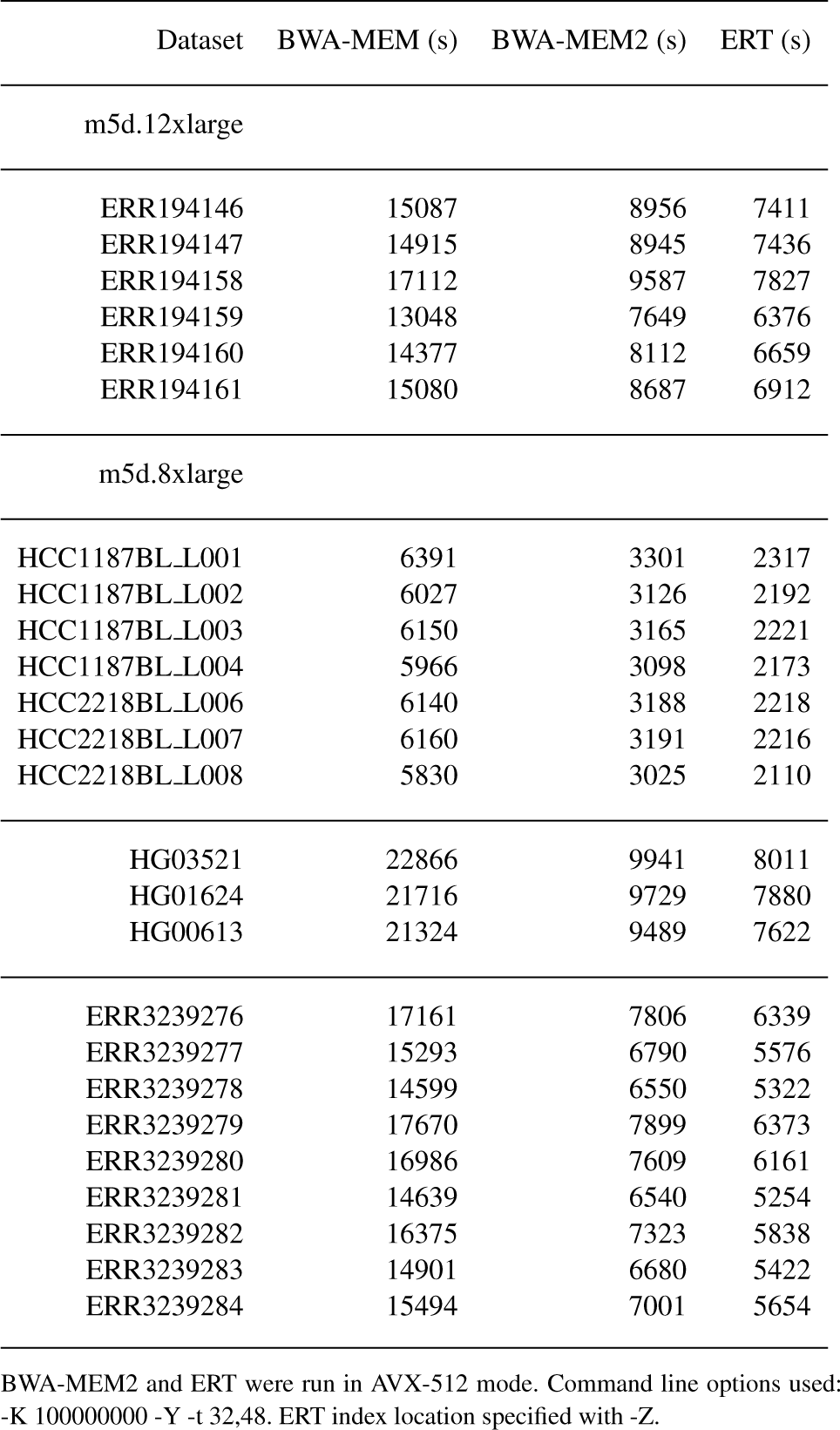
Wall-Clock Time for Overall Alignment

### 4.2 Seeding Performance

Figure 8 compares the performance of BWA-MEM, BWA-MEM2, CPU-ERT and FPGA-ERT for only the seeding step of read alignment. For this evaluation we use single-ended reads from the `ERR194147` dataset of Illumina Platinum Genomes. CPU and FPGA evaluations were performed on the `m5d.12xlarge` and `f1.4xlarge` instance respectively. It can be seen that CPU-ERT provides 2.1× higher seeding throughput than BWA-MEM2. FPGA-ERT can further improve the throughput of our CPU implementation by 1.6×.

**Fig. 8.**
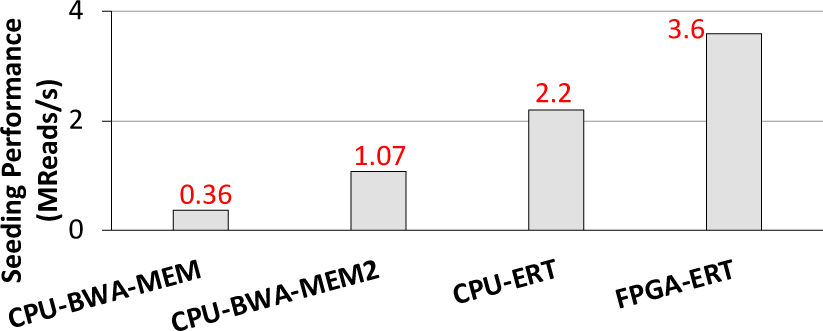
Seeding performance in Million reads/s.

### 4.3 Memory Access Breakdown

Figure 9 shows the breakdown of memory requests across different steps in ERT-based seeding for a set of 1 million reads sampled from `ERR194147`. By eliminating most of the redundant backward searches using a *zigzag* seeding approach (Section 3.4), we find that only 18% of the memory requests are to the index table. Nearly half of the memory requests in ERT seeding are due to tree traversal. Also only 5% of the requests are due to additional fetching of reference sequence needed to decompress leaf nodes, indicating that this step is not a performance bottleneck.

**Fig. 9.**
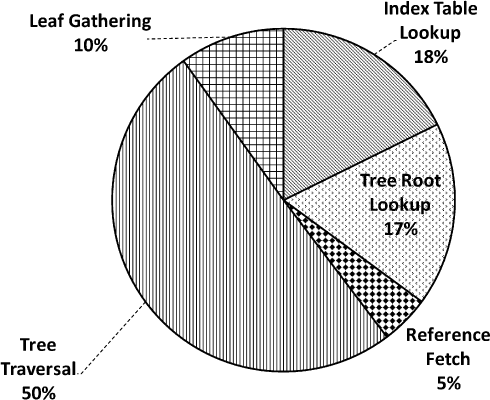
Distribution of memory requests across different steps of ERT-seeding

## 5 DISCUSSION

### Support for different read lengths

Like BWA-MEM and BWA-MEM2, ERT can also support different read lengths in the input dataset as long as the index has been built considering the maximum length of the read that can appear in the dataset. Maximum read length is used as a parameter to decide the maximum depth of the radix trees in ERT. For the human genome, we observe that the increase in the index size is small (1.3 GB) when maximum read length is changed to 251 from 151, with negligible impact on performance. This is because most leaf nodes appear close to the root. We therefore recommend that the maximum read length be set conservatively when building the ERT index.

### Equivalent output with BWA-MEM

Although ERT can be used standalone as an alternative for FMD-index, we chose to design ERT within the framework of existing aligners like BWA-MEM and BWA-MEM2 and take care to ensure the same output is produced. This was done with the goal of easing adoption and reducing validation effort.

### Index loading time

While index loading time was not significant using local and remote solid-state drives (SSD), the large index size of ERT like BWA-MEM2 can impact loading times in highly-loaded networked file systems with remote hard-disk drives. Our preliminary experiments suggest that `gzip` compression (∼19% reduction in ERT size) and parallel decompression can help alleviate some of the I/O costs of index loading. We leave this exploration to future work.

## 6 RELATED WORK

### CPU and GPU-based Seeding

FMD-index based seeding (Li, 2012; Ferragina and Manzini, 2000) involves many irregular memory accesses and has been found to be bottlenecked by LLC and TLB misses on CPUs (Chacón *et al.*, 2013; Zhang *et al.*, 2013). Prior work has explored reordering memory accesses (Zhang *et al.*, 2013) and performing *n*-character lookup on an *n*-step FMD-index (Chacón *et al.*, 2013) to improve the locality and data requirements of FMD-index based seeding. However, these implementations focus on exact-match search and do not natively support SMEM computation. There have also been efforts to improve the locality of the location operation using the FM-index (Cheng *et al.*, 2018) although these have not yet been integrated into read alignment software. Similar to ERT, recently released BWA-MEM2 (Vasimuddin *et al.*, 2019) also trades-off memory space for memory bandwidth, but by departing from the FMD-index and proposing a bandwidth-friendly data structure we further improve bandwidth efficiency while producing identical seeds. Data-parallel architectures such as GPUs have also been leveraged to accelerate FMD-index search (Chacón *et al.*, 2015) by virtue of their high memory bandwidth and memory level parallelism. However, ERT traversal is inherently not data-parallel and traversal patterns need to be optimized to reduce memory divergence in GPU’s SIMD units. We leave this exploration to future work.

While this work focuses on FMD-index based seeding, there exists a rich body of work that uses hash-tables for seeding (Ahmadi *et al.*, 2011; Kiełbasa *et al.*, 2011; Xin *et al.*, 2015) and have optimized its cache behavior (Hach *et al.*, 2010, 2014). Hash-based seeding coupled with filtration algorithms work well in mappers that produce large number of seeds for seed-extension.

### Seeding Accelerators

Seeding accelerators based on the FMD-index use custom bit-wise operations to traverse the index and improve memory parallelism (Wang *et al.*, 2018; Chang *et al.*, 2016; Cong *et al.*, 2018). However, since these implementations do not aim to reduce data requirements, they are limited by memory bandwidth. ERT takes a different approach by redesigning the seeding algorithm to reduce data requirements for seeding and unlocks significant potential for hardware acceleration.

### Radix Tree Applications

Suffix trees have been used to perform whole genome alignments in bioinformatics tools such as MUMmer v1.0 (Delcher *et al.*, 1999). However, because of their huge space requirements they have been less explored for performing read alignment against large genomes such as the human genome. Radix trees have been used to accelerate read mapping in a prior work (Tran and Chen, 2015), but to keep space requirements for the human genome tractable, they truncate the tree when a certain hit threshold is reached (*F* = 300). ERT overcomes this space limitation by eliminating long singleton-path tails in the tree and instead includes a pointer to the same string in the reference genome. This space-efficiency enables it to fit within the memory capacity on commodity servers (64 GB). Further when compared to conventional radix trees, ERT is highly customized for MEM-based seeding, optimized for spatial/temporal locality and can serve as a drop-in alternative for FMD-index seeding in read aligners such as BWA-MEM.

## 7 CONCLUSION

Seeding is an important time-consuming step in read alignment. Because of its small memory footprint, FMD-index is widely used for seeding in read alignment software such as BWA-MEM. However, FMD-index-based seeding is limited by memory bandwidth, primarily due to its character-by-character processing of reads. This paper demonstrates a hardware-software co-design approach that can be used to accelerate seeding by optimizing for memory bandwidth, rather than memory capacity. In particular: 1) we show that data-efficiency can be improved, at the cost of memory footprint by our technique for performing multi-character lookup and redesigning the seeding algorithm. 2) we show how to exploit spatial and temporal data reuse across operations through prefix-merging and k-mer reuse. We also highlight how data-efficiency improvements in seeding can expose significant acceleration potential by designing an FPGA prototype based on ERT. In summary, ERT improves read alignment throughput of BWA-MEM2 by 1.28×, while guaranteeing identical output to the original software.

## 8 SUPPLEMENTARY MATERIAL

### 8.1 Improving Temporal Locality of Seeding with K-mer Reuse

Figure 10 describes the steps to be performed to leverage k-mer reuse. While processing the forward extensions for a batch of *N* reads, we store each backward extension that must be computed in a k-mer metadata table implemented on-chip. Each backward extension entry is composed of: (1) k-mer starting from the backward extension point in the read, (2) the read ID in the batch, and (3) start position of backward extension in the read. Once all forward extensions have been completed for a batch of reads, we sort all entries in the on-chip memory, grouping each required backward extension by k-mer. We then proceed one k-mer at a time and compute all backward extensions associated with a k-mer sequentially. The first time a k-mer is encountered, we perform one index table lookup, as well as fetch of portion of the k-mer’s tree into an on-chip cache. Subsequent backward extensions then consult this cache during tree walking, skipping two otherwise mandatory DRAM accesses. If a backward extension needs data that does not exist in the tree cache, we fetch it from memory on-demand and store it in case future backward extensions require this data. K-mer reuse strictly decreases the number of radix trees fetched from DRAM–and reduces total bandwidth requirement–but adds the computational overhead of sorting the backward extensions by k-mer.

**Fig. 10.**
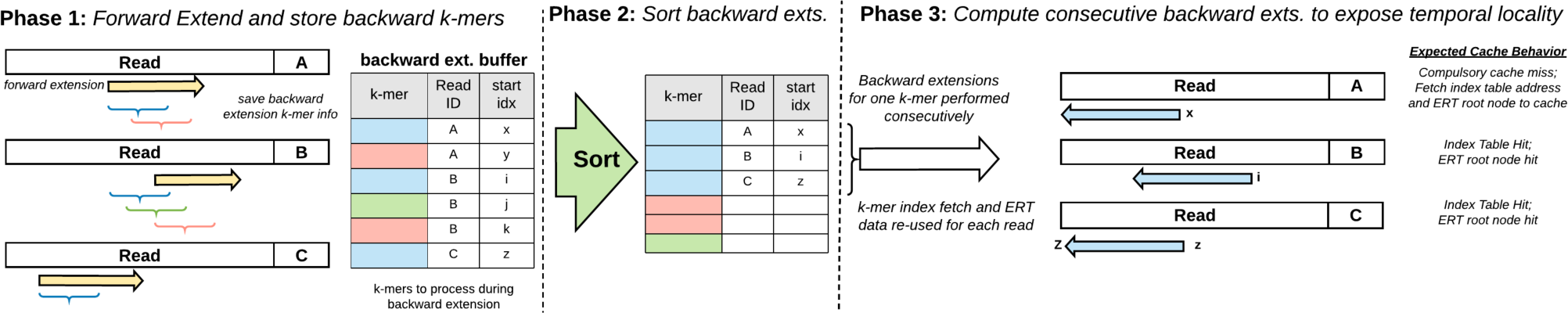
Phase 1) All unique k-mers required for backward extension are tracked during forward extension. Phase 2) k-mer information is sorted so all backward extension information for a single k-mer are in sequence. Phase 3) Backward extensions begin. The ERT for each k-mer only needs to be fetched once from DRAM, reducing bandwidth requirements.

### 8.2 Improving Spatial Locality of Seeding

Similar to (Kim *et al.*, 2010; Chilimbi *et al.*, 1999), we adopt a tiled layout for the nodes of the radix tree to improve spatial locality of accesses (Figure 11). In this layout, subtrees of nodes that are likely to be accessed at the same time are clustered together into a single cache block- or a DRAM page-sized tile. Compared to breadth-first or depth-first layout of nodes, the tiled layout guarantees at least *log*_4_(*n* + 1) nodes accesses per tile, where *n* is the number of nodes in the tile. With this optimization, ERT traverses ∼3 nodes on average per 64 B, utilizing 50% of the data it fetches from memory.

**Fig. 11.**
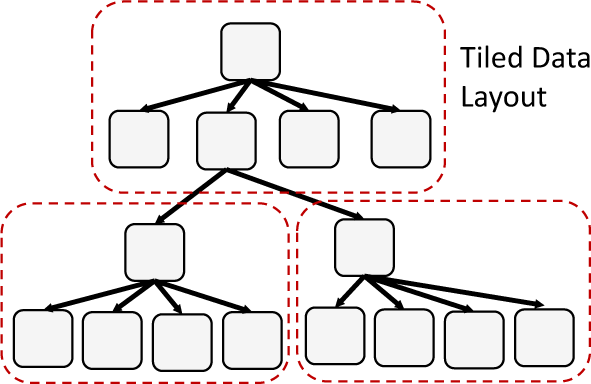
Cache-friendly tiled data-layout for ERT.

